# Evolving synergetic interactions

**DOI:** 10.1101/028357

**Authors:** Bin Wu, Jordi Arranz, Jinming Du, Da Zhou, Arne Traulsen

## Abstract

The outcome of a major evolutionary transition is the aggregation of independent entities into a new synergetic level of organisation. Classical models involve either pairwise interactions between individuals or a linear superposition of these interactions. However, major evolutionary transitions display synergetic effects: their outcome is not just the sum of its parts. Multiplayer games can display such synergies, as their payoff can be different from the sum of any collection of two-player interactions. Assuming that all interactions start from pairs, how can synergetic multiplayer games emerge from simpler pairwise interaction? Here, we present a mathematical model that captures the transition from pairwise interactions to synergetic multiplayer ones. We assume that different social groups have different breaking rates. We show that non-uniform breaking rates do foster the emergence of synergy, even though individuals always interact in pairs. Our work sheds new light on the mechanisms underlying a major evolutionary transition.

## 1 Introduction

Major evolutionary transitions share cooperation as a common theme: simple units aggregate to form a new level of organisation in which individuals benefit others bearing a cost to themselves (1, 2). However, from a Darwinian perspective, cooperation is difficult to explain, as natural selection promotes selfishness rather than cooperation (3–7). The evolution of cooperation has often been approached through the lens of simple two-player games that depict social dilemmas (8–10). The study of games such as the Prisoner’s Dilemma, or alternative situations such as the Stag-Hunt game (11), have provided insightful views on which mechanisms are likely to promote cooperation — e.g., spatial reciprocity, direct reciprocity, indirect reciprocity, kin selection and group selection (12–14). However, the simplicity of two-player games is a double-edged sword, as these pairwise games may fail to capture the intricacies of complex interactions in real social and biological systems. Evolutionary transitions typically involve multiple interaction partners at the same time rather than a collection of pairwise interactions. For instance, when cells interact to form a multicellular organism, a superposition of pairwise interactions is insufficient to capture the intricacies of the complex organism. This is because an interaction among all the cells is not just a sum of pairwise interactions. Synergetic interactions — the whole being more than the sum of its parts — may be necessary to allow a high level of selection unit to emerge. Synergetic interactions could then pave the way for the emergence of complex phenomena such as division of labour or multicellularity. Therefore, understanding how synergetic interactions emerge is an important part of our understanding of evolutionary transitions.

General multiplayer games, which cannot be decomposed into pairwise interactions, can represent such synergy effects. They can display broader and richer dynamics than their traditional two-player counterparts (15–18). In particular, multiplayer games can exhibit payoff non-linearities and can thus account for the synergetic effects that are intrinsic to major evolutionary transitions. Although the emergence of synergetic interactions among multiple players is key to all major evolutionary transitions as aforementioned (19), we lack fundamental understanding on how such complex synergetic interactions occur in the first place. Here, we present a mathematical model that captures the emergence of synergetic multiplayer games from simple pairwise interactions.

## 2 Results

### 2.1 Model description

We consider a structured population of *N* individuals, assorted into *l* sets (20), each consisting of *m* individuals. Individuals can have a different number of set memberships and play one of two strategies, *A* or *B*. An individual accumulates the payoff through interactions within all the sets it belongs to. These interactions are always pairwise and the payoff depends on the set configuration, i.e., the number of individuals playing *A* and *B* in the set. At every time-step, either the strategy of an individual or the set structure is updated. With probability *w*, the strategy of an individual is updated. Two individuals are randomly chosen and one imitates the other’s strategy with a probability that increases with the payoff difference. Individuals with higher payoffs are more likely to be imitated (21, 22). With probability 1 – *w*, a set is randomly chosen. This set may break up with probability *k_i_*, — where *i* is the number of strategy *A* individuals in the set, ranging from 0 to *m*. As a consequence, *k_i_* determines the fragility of a set, which in turn, depends on the set composition (23). If the set breaks, a randomly chosen individual — which belongs to at least one other set — is expelled. In order to keep the size of the set constant, another random individual is then incorporated into the set (Fig.1).

**Figure 1:**
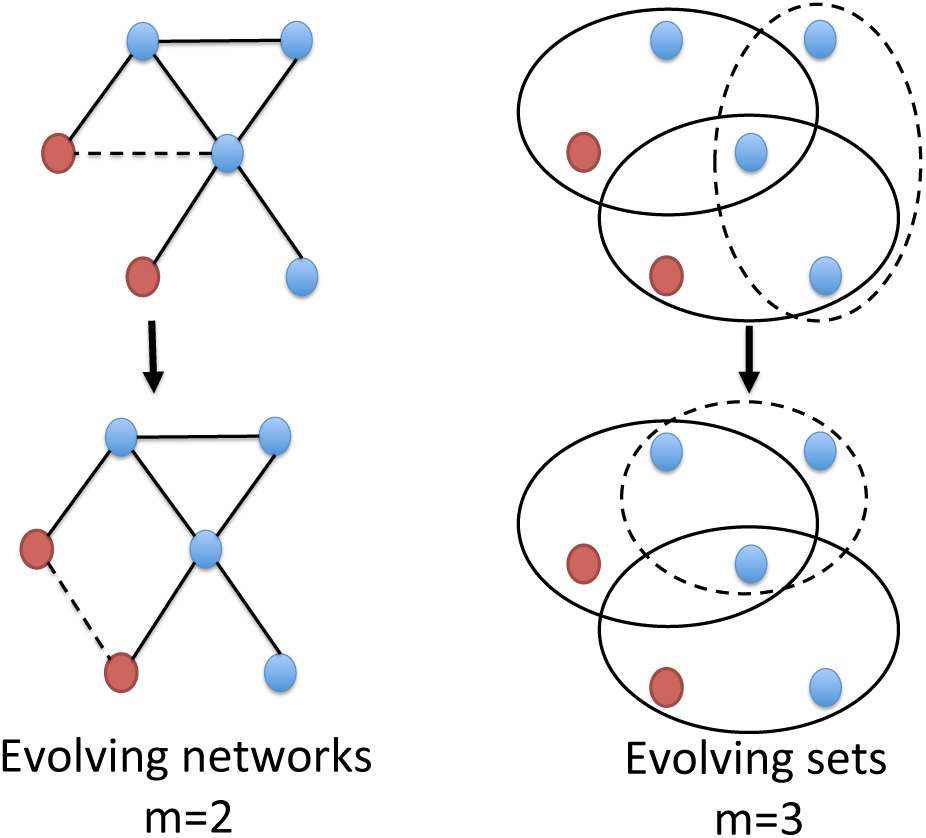
Set dynamics for sets of size *m* = 2 (left column) and *m* = 3 (right column). Blue and red dots represent strategies *A* and *B* respectively. When *m* = 2 (left column) “sets” are actually “links” and the overall structure is a network (26). In this case, interactions are strictly pairwise, hence there is no synergetic effect in the payoffs. The right column shows the case with *m* = 3, which is the minimum set size that illustrates the emergence of synergetic interactions. With probability 1 − *w* a set is selected at random (dashed lines). This set breaks up with probability *k_i_*, where *i* is the number of strategy *A* individuals in the set. If the set breaks, a randomly chosen individual is expelled. In order to keep the size of the set constant, another random individual is incorporated into the updated set (dashed lines).

Although our model is simple, it captures two fundamental aspects. First, the set structure mimics the social interactions which is an intrisic characteristic of biological systems and human societies. For instance, the set size could be based on the diffusion rate of the public goods secreted by cooperative cells (24, 25). The overall structure also allows to consider an organisation of arbitrary size, from a small family to a large assembly. Second, individuals only interact in pairs and payoffs are additive. In this case, the payoff of an individual is nothing but sum of the corresponding pairwise interactions. Thus, there are no imposed synergetic effects via the payoff accumulation process. Instead, it can only emerge from the dynamics of the population structure.

#### Pairwise games between two strategies

If the size of the sets is two, *m* = 2, the population structure is equivalent to a network, where a “set” becomes a “link” (Fig. 1). In this case, the accumulated payoffs can still be captured by a pairwise interaction, hence there is no synergetic multiplayer interactions (26, 27). Although our analytical framework is general enough to allow the study of any set size, we focus on the case where *m* = 3, that is, when the sets contain three individuals. Let us assume that for two individuals playing strategy *A* (*B*), each one obtains a payoff of *a*(*d*). Similarly, for two individuals playing different strategies, the individual using strategy *A* obtains the payoff *b* and the individual using strategy *B* obtains the payoff *c*. Given that there are three individuals in every set, the payoff of an individual within a set is determined by two pairwise interactions. Therefore, the payoff of an individual playing strategy *A* (*B*) in a set with *j* other individuals playing *A* is given by *a_j_ = aj* + *b*(2 − j) (*b_j_ = cj* + *d*(2 *− j*)). Note that for *m* = 3, *j* = 0, 1, 2. Given that the breaking probability of a specific set may depend on the number of *A* individuals within the set, the set dynamics allows for non-uniform breaking probabilities across the sets.

To demonstrate that our simple model can indeed capture the emergence of synergy, we consider two aspects: the accumulated payoff of both types and the evolutionary dynamics of the two strategies. We find that non-uniform breaking probabilities across the sets foster the emergence of synergetic multiplayer interactions. In other words, when the fragilities of the sets are non-uniform we find that (i) the expected accumulated payoff of both strategies is consistent with the one of a typical multiplayer game that cannot be decomposed into a pairwise game, and (ii) the evolutionary dynamics of the strategies exhibit two internal equilibria of selection, which is impossible in a two-player game.

The calculation of the average accumulated payoff in the general case is challenging, even though the model is simple. We overcome this problem by assuming that the probability with which the strategy is updated is small, *w* ≪ 1. Asa consequence, the set structure can reach its stationary state—which determines the accumulated payoffs—before a strategy update occurs. Importantly, the average accumulated payoffs for both strategies are consistent with the payoffs of the following 3-player game in a well mixed population, up to a positive rescaling factor (see SI Appendix, Section 2.1):

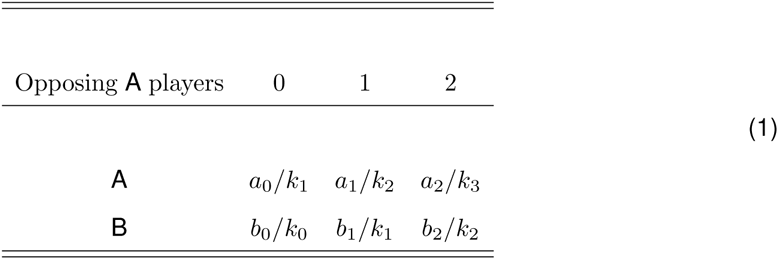

Here *a_i_/k_i+1_* is the payoff for an individual using strategy *A* when it meets *i* opponents using strategy *A*. Equivalently, *b_i_/k_i_* is the payoff for an individual using strategy *B* when it meets *i* opponents using strategy *A*. The payoff table in Eq.(1) has two important features. First, the derived multiplayer game is of the same size as that of the set. Second, the payoff entries are proportional to the product of the accumulated payoff in a set and its lifetime.

The evolutionary outcome of both strategies can be predicted by the replicator equation for a large class of microscopic imitation rules, if the population size is sufficiently large (see SI Appendix, Section 2.2). The replicator equation is given by

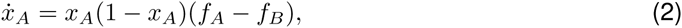

where

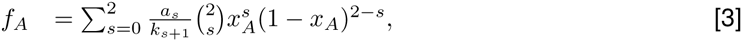

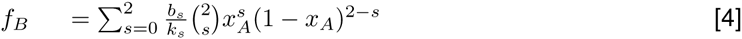

are the payoffs for strategy *A* and B of the 3-player game based on Eq. (1), and *x_A_* is the fraction of individuals using strategy *A*. In other words, the dynamics of the pairwise game under active set dynamics can be captured by a multiplayer game in a well-mixed population. The internal equilibria of this equation are the roots of the equation *f_A_ − f_B_* = 0. Based on the initial fraction of individuals using strategy *A*, these equilibria determine where an infinite population would end up (28).

The above results on the accumulated payoffs and the evolutionary dynamics of strategies hold for any set fragilities (*k*_0_, *k*_1_, *k*_2_, *k*_3_). In the following, we apply these results to homogenous and heterogeneous set fragilities to address when and how synergetic interactions emerge.

Whenever the fragility of the sets is homogenous, *k*_0_ = *k*_1_ = *k*_2_ = *k*_3_, Eq. (1) is identical to the one of the original pairwise game, even though it is a 3-player game (see SI Appendix, Section 2.2). Therefore there is no synergy effect in the payoffs. Given that the replicator equation is equivalent to the one of the pairwise game, there is at most one internal equilibrium with the same position and stability. The upper panel of Fig. 2 shows the agreement between the analytical approximation and a simulation of the full model.

**Figure 2:**
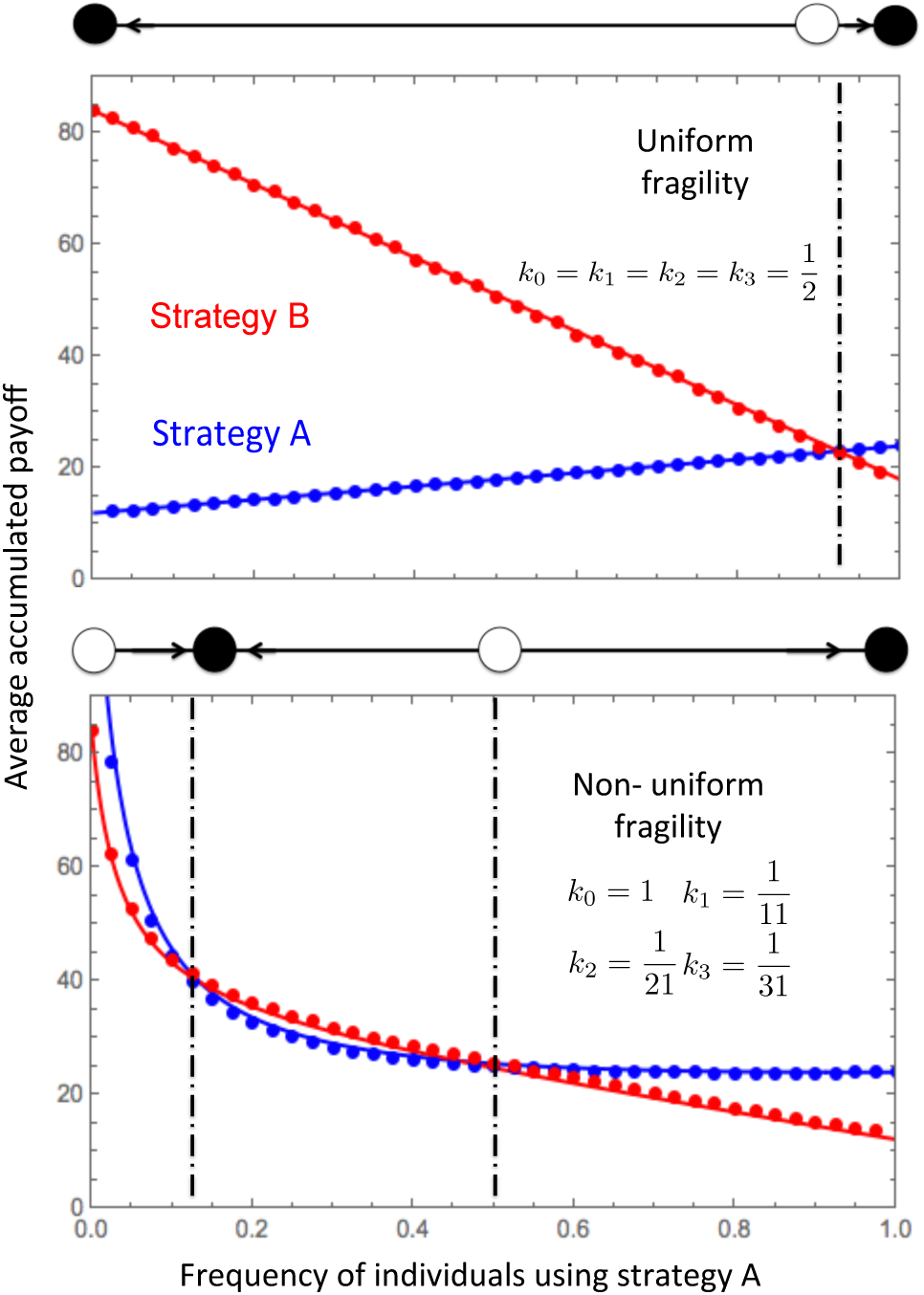
Accumulated payoffs for the Stag-Hunt game and the equilibria of the selection gradient for two fragility scenarios. The full/empty circles on top of each of the two panels show the analytical approximation for the stable/unstable equilibria of the replicator dynamics. Top: For uniform breaking rates, the accumulated payoffs for both strategies match the ones of a pairwise game. Thus, the payoffs change linearly with the fraction of *A* individuals and the replicator equation predicts just the single internal equilibrium of the pairwise game. Bottom: When the breaking rates depends on the set configuration, the payoffs for both strategies become non-linear. The more *A* individuals a set has, the less likely it breaks up. The payoffs for both strategies have two intersections, which lead to two internal equilibria. This illustrates that non-uniform interactions can lead to the emergence of synergetic interactions. There is a perfect agreement between simulations (dots) and analytical approximation (lines). (Parameters: Stag-Hunt game with *a* = 2, *b* = 1, *c* = 1.5 and *d* = 7. Population size, *N* = 500, number of sets, *l* = 1000, probability of a strategy update, *w* = 10^−3^. Selection intensity, *β* = 0.1. See Methods for the simulation details).

However, when fragilities are not homogeneous across the sets, Eq.(1) becomes a 3-player game, which cannot be decomposed into additive pairwise interactions (lower panel of Fig. 2). In this case, the payoff of an individual interacting with two opponents is not equal to the sum of the two pairwise interactions (Fig. 3). Consequently, the presence of non-uniform set breaking probabilities generates synergetic payoffs. Synergy emerges exclusively as a result of the evolutionary dynamics of the set structured population. In addition to this, the replicator equation has two internal equilibria, which is not possible in pairwise interactions (see lower panel of Fig. 2). Static random networks display similar effects (29). A necessary condition for the emergence of two equilibria is that the sign of the effective payoff difference *a_i_/k_i+1_ − b_i_/k_i_* changes twice with the increase of the number of opponents using strategy *A*, *i* (30, 31). A more detailed analysis on the conditions that lead to two internal equilibria can be found in SI Appendix, Section 2.2. If one of the two equilibria is stable, the other has to be unstable. Given this, our model can explain both the maintenance of biodiversity and phenotypic dominance within the same framework.

**Figure 3:**
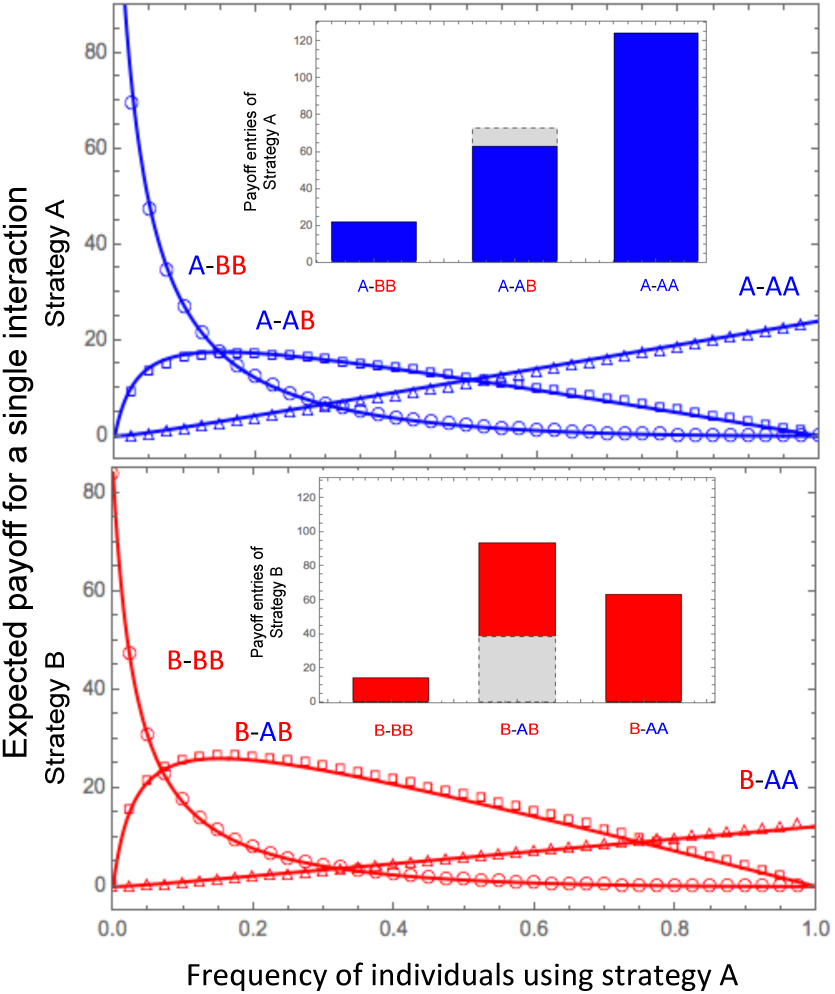
Expected payoffs for individuals using strategies *A* (upper panel) and *B* (lower panel) in groups of different compositions. The two insets within each plot show the effective payoff entries of the emergent 3-player game Eq. (1) for a frequency of 0.5. Main panels: Theoretical predictions for the payoff within the three set configurations of each individual, Eqs. (3) and (4) (lines) agree well the accumulated payoff obtained by simulation (symbols). This in turn proves that the synergetic 3-player game is intrinsically captured by Eqs. (3) and (4) even term by term. Upper inset: Payoffs for equal abundance of both strategies. An individual with strategy *A* gains less if it interacts in a set which has 1 individual with strategy *A* than it were in the synergy free case (grey, dashed). Lower inset: An individual with strategy *B* gains much more if it interacts in a set which has 1 individual with strategy *A* than it were in the synergy free case (grey, dashed). Here the payoffs in the synergy free cases are the average values of the two payoff entries for the focal individual interacting with 0 and 2 *A* individuals. Thus, the interaction can no longer be decomposed into multiple pairwise interactions, which is how every individual obtains its payoff microscopically. (Parameters are the same as that in the lower panel of Fig. 2. The inner panels are obtained by simulation via setting the frequency of individuals using strategy *A* to be one half. See Methods for the simulation details.)

#### Pairwise games between *n* strategies

The model can be extended to account for an arbitrary number of strategies, *n*, instead of only two. In the pairwise interactions with n strategies or an *n* × *n* game, the non-uniform breaking probabilities also generate synergetic multiplayer interactions. Although the analytical calculations are more intricate due to the increased number of set configurations, we find that the payoff matrix of the emergent multiplayer game is consistent with the one of an *n*-strategy *m*-player game (SI Appendix, Section 3.1). Interestingly, the intuition behind these payoff entries is similar to the ones of the two-strategy case, as they still represent the product of the additive payoffs via pairwise interactions and the duration of that set. In addition to this, the *n*-strategy *m*-player game has, at most, (*m* − 1)^n−1^ isolated internal equilibria, where as the original *n × n* pairwise game has at most one equilibrium (SI Appendix, Section 3.2). The dynamics in our model are thus rich enough to capture complex phenomena exhibited by social and biological systems.

## 3 Discussion

Synergy refers to the idea that the whole is greater than the sum of its parts. Interestingly, synergy is identical to “cooperation” in ancient Greek. Synergy can be observed in a plethora of different contexts such as in genes (32), microbial populations (33), and even social and economic systems. From an evolutionary perspective, synergy is a cornerstone of all major evolutionary transitions. These evolutionary milestones involve the aggregation of simple units into a new entity which becomes a higher-level Darwinian unit of selection (1). With this in mind, we present a minimalistic model that shows how synergy can actually emerge. Our model allows to treat the emergence of synergetic interactions from simple additive pairwise ones analytically. We assume that the strategy of the individuals and the set structure evolve in time. The results prove that non-uniform set breaking rates, which depend exclusively on the configuration of these sets, lead to payoff non-linearities. These are consistent with the dynamics of a multiplayer game, even though individuals always play a two-player game and no group selection effects are present. These findings rely on two conditions: i) sets must contain more than two individuals, and ii) the breaking rates must depend on the configuration of the sets and, hence be non-uniform. As a consequence, our model may be useful as a starting point for the investigation of the evolution of more complex phenomena, e.g., synergetic interactions within the group. It shows how the aggregation of individuals can lead to complex interactions that cannot be disentangled into simpler interactions.

## Methods

*The Fermi updating rule.* We use the Fermi update rule, given by the following algorithm:

i. Randomly select an individual, a* and denote its payoff as *π_a_\**;
ii. Randomly select another individual, b*, among all the individuals in the sets individual *a*\* is in and denote the payoff of b* as *-π_b*_*;
iii. *a*\*switches to the strategy of *b*\* with probability (1 + exp[−*β*(π*_b_*_*_ − π*_a_*_*_)])^−1^.

*Accumulated payoffs.* Each data point is the average of 100 independent realisations. Every realisation takes 10^6^ generations. In each realisation, for the first 10^4^ generations, only set dynamics occur. For the rest generations, at every step, with probability *w* = 10^−3^ we compute the average accumulated payoff of each strategy. Otherwise, with probability 1 − *w*, set dynamics happens. At the end of each realisation, we compute the mean value of all the average accumulated payoff. *Selection gradient.* Each data point is the average of 100 independent realisations. Every realisation takes 10^7^ generation. For the first 10^7^ generations of each realisation, only set dynamics occur. After that, with a probability of *w* = 10^−3^ two individuals are chosen randomly from the entire population. The first individual is the “focal” one which is the one that may imitate the strategy of the second one based on the Fermi rule. We keep track of the transition without implementing them. We denote *y* and *z* as the number of times that an individual playing strategy *A* and *B* changes its strategy. 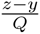 is the estimator of the selection gradient 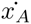, where *Q* is the number of strategy updating events in this realisation.

*Accumulated group payoffs.* Each data point is the average of 100 independent realisations. Every realisation takes 10^6^ generations. In each realisation, for the first 10^4^ generations, only set dynamics occur. For the rest generations, at every step, with probability w = 10^−3^ we compute the average accumulated payoff of each strategy induced by the set with 0, 1 and 2 strategy A opponents, respectively. Otherwise, with probability 1 − *w*, set dynamics happens. At the end of each realisation, we compute the mean value of all the average accumulated payoff.

## Acknowledgements

We thank Chaitanya Gokhale for inspiring discussions.

## Supplementary Information of “Evolving synergetic interactions”

### 1 Evolutionary dynamics in a set structured population

Consider a set structured population of fixed size *N* where individuals engage in a pairwise game. The population is divided into a constant number of sets, *l*, each of them of constant size *m*. Individuals can belong to different sets.

At each time step, we either update the strategy of an individual — with probability *w* — or the structure of the population, with probability 1 – *w*.

A strategy update involves two randomly selected individuals, say Alice and Bob. Alice imitates Bob’s strategy with a probability depending on the difference of their payoffs, i.e., the imitation rule (1). The payoff of an individual is calculated as the sum of all the pairwise interactions through all the sets it belongs to. For instance, if Alice belongs to two sets, her payoff is the sum of the payoff in the first set, plus the payoff in the second one. Given that every set consists of m individuals, the payoff of an individual in a set is the sum of *m* – 1 pairwise interactions. On the other hand, when set dynamics occur, a set is randomly selected. This set may break with a probability which depends on the set composition. In particular, if there are only two strategies, the set composition is the number of individuals within the set playing a specific strategy. If the set is broken, a random individual within the set is expelled, provided it is in at least one other set. In order to keep the size of the set constant, another random individual is added to the focal set.

We start with the simplest pairwise games with two strategies, *A* and *B*. The 2 × 2 payoff matrix is given by (*a_ij_*)_2×2_, where *a_ij_* is the payoff of an individual playing strategy *i* with an opponent playing strategy *j*, where *i,j ∈* {*A, B*}. We find that the average accumulated payoff for each strategy is consistent with the one of a 2-strategy, *m*-player game up to a positive rescaling factor. When the breaking probability of a set is uniform across all kinds of sets, the payoff of the *m*-player game is equivalent to that of a sum of *m* – 1 pairwise games. However, we notice that whenever the sets have different breaking probabilities that depend on the set composition, intrinsic multiplayer interactions emerge. In this case, the accumulated payoff of both strategies cannot be decomposed into collections of pairwise games anymore. In other words, synergetic effects in payoff can emerge from simple pairwise interactions. Based on accumulated payoffs, we further obtain the replicator equation of the *m*-player game to determine the evolutionary fate of each strategy. In addition to this, the replicator equation shows up to *m* – 1 internal equilibria. In contrast, for pairwise interaction there is at most one such equilibrium. These results are obtained under the assumption of fast set dynamics — very few strategy updates occurs before the population structure has reached the stationary state — and a large population size.

We generalise the above results for cases where the number of strategies, *n*, is greater than two. In this case the payoff matrix is given by (*a_ij_*)*_n × n_*, where *i,j ∈* {1,2,&, *n*}. In this technically somewhat more challenging case, we find similar results:

i. The average accumulated payoff of each strategy is of the form of an *n*-strategy, *m*-player game up to a positive rescaling factor.
ii. When the set breaking probabilities are uniform, the payoff of the *n*-strategy *m*-player game is still consistent with the sum of the *m* – 1 pairwise games.
iii. Non-uniform set breaking probabilities foster the emergence of multiplayer interactions, which cannot be decomposed into a collection of pairwise games.
iv. The replicator equation of the *n*-strategy *m*-player captures evolutionary dynamics of the strategies and displays, at most, (*n* − 1)*^m^*^−1^ internal equilibria, whereas pairwise *n × n* games display, at most, a single equilibrium (2).

### 2 Games with two strategies

#### 2.1 Accumulated payoffs

Initially, i.e. at time *t* = 0, we call the *l* sets 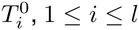. For the first time step in the set evolution, *t* = 1, we denote the selected set as 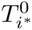. If this set is broken and transforms to another set, we denote the transformed set as 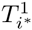 otherwise the set is not broken and we let 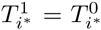. For the other sets which are not selected, we denote 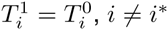. Recursively, we define 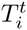 for t ≥ 0 and 1 ≤ *i* ≤ *l*.

Let 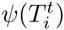 be the number of strategy *A* individuals in set 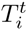, thus 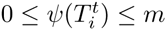. For each set *i*, the dynamics of 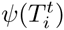 is a Markov chain in state space {0,1,2 … m} with the transition matrix

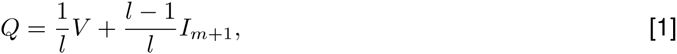

where *I_m+_*_1_ is the identity matrix of size *m* + 1 and *V* is the transition matrix when set *i* is selected.

Hence, *V* = (*V_ij_*)_(_*_m_*_+1)×(_*_m_*_+1)_ (0 ≤ *i,j ≤ m*) is a tridiagonal matrix given by

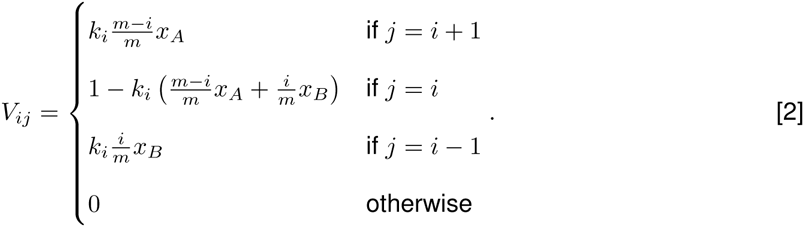

Here *x_A_* and *x_B_* = 1 − *x_A_* are the fractions of strategy *A* and *B* in the population, *k_i_* is the breaking probability of a set consisting of *i* individuals playing strategy *A* and *m* − *i* individuals playing strategy *B*.

When *x_A_* > 0 and *x_B_* > 0, the matrix *Q* is irreducible and aperiodic and there is a unique stationary distribution *y* = (*y*_0_,*y*_1;_ … *,y_m_*) of *Q* determined by *yQ* = *y.* Taking Eq. (1) into *yQ* = *y* leads to *yV = y*. This leads to the stationary distribution

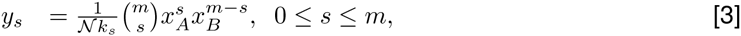

where 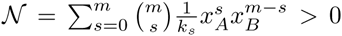 0 is a normalisation factor. The stationary distribution also represents the proportion of each type of set among all the sets in the stationary regime.

When set dynamics are fast, the imitation event happens rarely enough to allow the population structure to reach the stationary state before a single imitation event occurs. In this case, the stationary regime of the set dynamics determines the average payoff of both strategies. Here the accumulated payoff of strategy *A* is given by

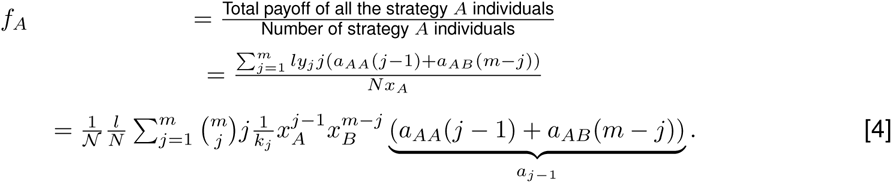

As 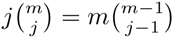, Eq. (4) can be written as

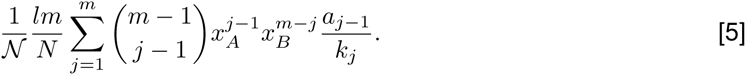

With s = *j −* 1, Eq(5) becomes

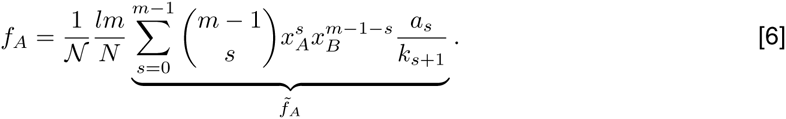

Similarly, we have

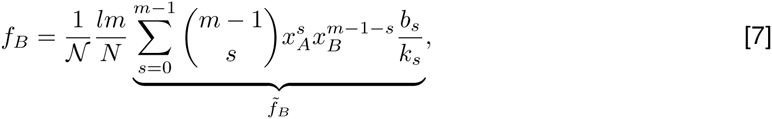

where *b_s_* = *a_BA_s + a_BB_*(*m – 1 – s*) is the accumulated payoff of an individual using strategy *B* gets in a set consisting of *s* individuals using strategy *A*.

Besides the common positive rescaling factor 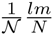, the average accumulated payoffs for both strategies are 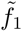 and 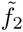. Interestingly 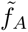 and 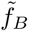 are exactly the payoff of an *m*-player game in a well-mixed population,

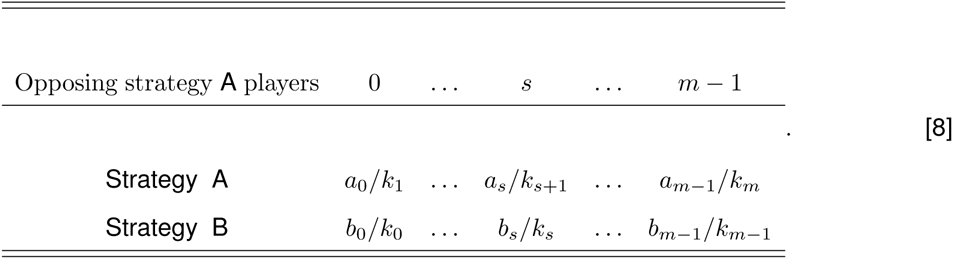

Here *a_s_/k_s+_*_1_ (*b_s_/k_s_*) is the payoff of an individual using strategy *A(B)* obtains when it meets s strategy *A* opponents. This payoff table has two remarkable features: First, the number of players in the multiplayer game and the set size are equal. Second, the payoff entries of the multiplayer game are the product of the linear collective payoff via pairwise interactions in a set and the duration of the corresponding set. Taking *a*_0_*/k*_1_ as an example, *a*_0_*/k*_1_ is the payoff of an individual playing strategy A when he interacts with 0 strategy *A* opponents, or *m* – 1 strategy *B* individuals. However, within a set, the accumulated payoff of an individual playing strategy *A* is given by *a_0_* = (*m − 1) a*_12_ and results from the *m* – 1 pairwise interactions in the set where 1/*k*_1_ is the duration of the set. In other words, *a*_0_*/k*_1_ is the accumulated payoff rescaled with the interaction time.

When *m* = 2, the population strcture is equivalent to a network and the sets represent links. In this case, the effective payoff table in Eq. (8) is still a pairwise game. This transformation can alter the effective payoff of both strategies, however it cannot lead to synergetic effect in payoffs as there is only one pairwise interaction in the transformed table. When *m* = 3, the emergent payoff table is consistent with a 3-player game. On one hand, we can take the payoff entries as the synergetic payoff of two individuals. On the other hand, we have the additive payoffs via the two pairwise interactions in that set. A comparison between these two payoffs can facilitate us to study when “the whole is better than the sum of its parts”, i.e., the synergetic payoff is better off than that derived by two pairwise interactions.

When the set breaking probabilities are uniform — i.e., *k*, is constant —, we find — by Eq. (6) and Eq. (7)—that the accumulated payoffs for strategy *A* and *B* are *f_A_* = (*m* – 1)(*a*_11_*x*_A_+*a*_12_*x*_B_) and *f_B_* = (*m* – 1)(*a*_21_*x*_A_ + *a*_22_*x*_B_). That is to say that the emergent payoff is equivalent to the sum of the corresponding *m* – 1 pairwise game and therefore there is no synergy. However, when the set breaking probabilities are non-uniform, intrinsic multiple player games emerge. In this case the “whole” is different from the sum of its parts.

#### 2.2 Evolutionary dynamics of strategies

In large well mixed populations, the evolutionary dynamics of strategies based on the imitation rule can be approximated by the Langevin equation (3)

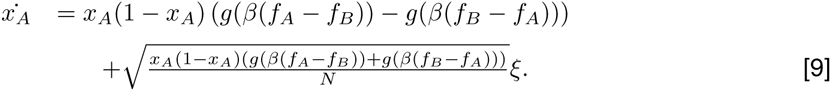

Here ξ is the white Gaussian noise, *f_A_* (Eq. 6) and *f_B_* (Eq. 7) are the average accumulated payoffs for strategy *A* and *B*, respectively. In addition, *g* is the imitation function capturing the likelyhood of the focal individual to adopt the strategy of the opponent’s and *β* is the selection intensity (4). Throughout, *g′* is positive, implying that individuals are likely to adopt the strategy of individuals with high payoffs. In particular, the Fermi update rule is an imitation update rule with the imitation function *g*(*x*) = [1 + exp(−*x*)]^−1^.

For large population size *N*, the stochastic term vanishes and we obtain

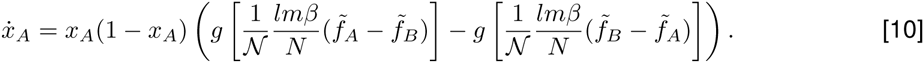

Note that 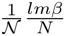 is always positive and *g*′ > 0, the equilibria of this equation are the same as that of the following replicator equation of the multiplayer game Eq. (8) in position and stability

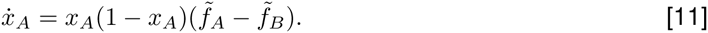

Therefore, the evolution of a pairwise game on the evolving set structured population is captured by an *m*-player game in a well mixed population. Under uniform breaking probabilities, the replicator equation Eq. (11) is consistent with the one of the pairwise game in well-mixed population. At most one internal equilibrium can arise in this case. Under non-uniform breaking probabilities, however, Eq. (11) can exhibit up *m* – 1 internal equilibria.

For any set size *m*, the internal roots of the replicator equation are determined by the roots of the following Bernstein polynomial (5)

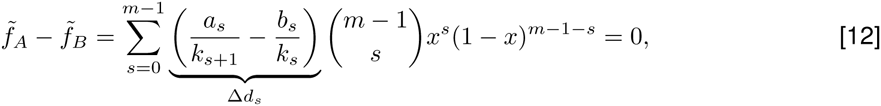

where *x* ∈ (0,1). By the variation diminishing property (6) we know that the number of the internal roots is equal to the number of sign changes of (Δ*d*_0_, Δ*d*_1&_, Δ*d_m_*_−1_), or less by an even number. In particular, when there is only one sign change in the sequence (Δ*d*_0_, Δ*d*_1_ …, Δ*d_m_*_−1_), there is exactly one internal equilibrium.

When the set size *m* is 3, there can be, at most, two internal equilibria (Fig. 1). A necessary condition for the existence of two equilibria is either Δ*d*_0_ > 0 Δ*d*_1_ < 0 and Δ*d*_2_ > 0, or Δ*d*_0_ < 0 Δ*d*_1_ > 0 and Δ*d*_2_ < 0. In both cases, the sign of the coefficient changes twice. The variation diminishing property tells us that there can be either two or no internal equilibria. Since Δ*d*_0_ > 0 Δ*d*_1_ < 0 and Δ*d*_2_ > 0 is equivalent to Δ*d*_0_ < 0 Δ*d*_1_ > 0 and Δ*d*_2_ < 0 by exchanging the name of the two strategies. We focus on Δ*d*_0_ < 0 Δ*d*_1_ > 0 and Δ*d*_2_ < 0. In this case, the Bernstein polynomial Eq. (12) is negative at *x* = 0 and 1. Thus, the existence of two internal equilibria is equivalent to that the maximum of the Bernstein polynomial in (0,1) has to be positive. In the present case, the Bernstein polynomial is quadratic with a maximum at *x*\* = (Δ*d*_1_ − Δ*d*_2_)/((Δ*d*_1_ − Δ*d*_2_) + (Δ*d*_1_ − Δ*d*_0_)) ∈(0.1). Therefore, the Bernstein polynomial is positive at *x\** if

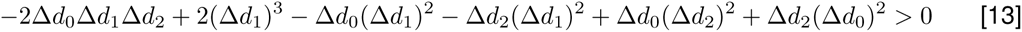

To sum up, there are two internal equilibria if and only if either of the two conditions holds

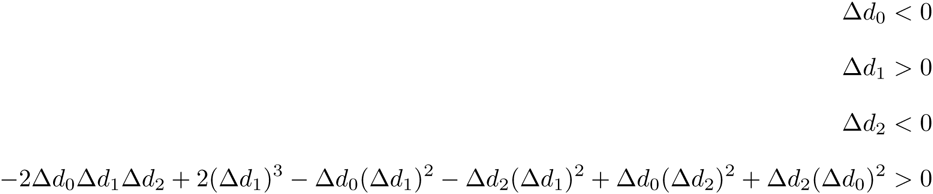

or

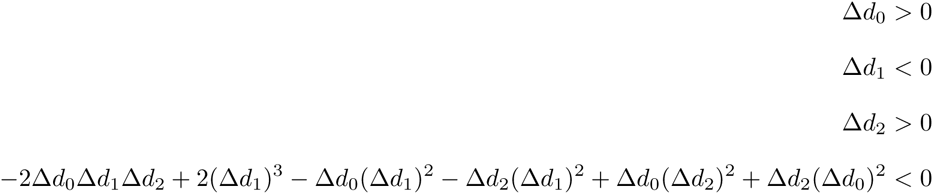

### 3 Games with *n* strategies

In the above section we assumed that each individual plays a pairwise game with its opponent. In addition, every individual can choose only between 2 strategies. In this section, we allow individuals to choose any number of strategies and thus generalise our analysis to *n* strategies. In this case, the pairwise interaction becomes an *n × n* game. We show that the previous results also hold for *n* strategies when the set dynamics are fast enough as, (i) the accumulated payoff for any strategy is an *m*-player game and (ii) the evolutionary dynamics of strategies can be captured by the replicator equation of the *n*-strategy *m*-player game.

#### 3.1 Accumulated payoffs

Similar to the 2-strategy case, the breaking probability of a set depends exclusively on its strategy composition. Let us denote 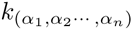 as the breaking probability of a set where *α_s_* is the number of strategy-*s* individuals in the focal set and *α_s_ ≥* 0 and 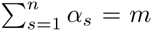 indicates that the set consists exactly of *m* individuals.

We start by randomly choosing one of the *l* sets, namely *i*. Then we define a sequence of sets 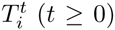. Here the set 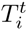 evolves into 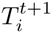. The type of the set 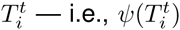 — is a Markov chain whose states are given by the possible set configurations. These set configurations can be denoted as the simplex

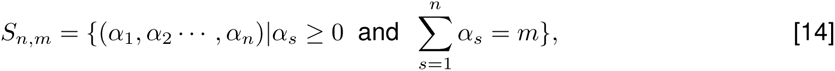

where *α_s_* is the number of strategy *s* individuals in the corresponding set. The transition matrix of this Markov chain is given by

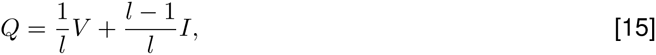

where *I* is the identity matrix of size *|S_n_,_m_|*. Here *|S_n_,_m_|* is the cardinal number of set *S_n_,_m_*. *V* is the transition matrix conditioned on the fact that the set *i* is selected. By the updating rule of the sets, two subsequent sets 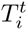 and 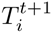 have at least m – 1 individuals in common. Thus the transition is impossible between two states (*α*_1_*,α_2_* …*,α_n_*) and 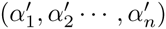, unless either of the following two cases holds.

- There exist two different strategies *s*_1_ and *s*_2_ such that 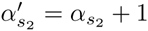 and 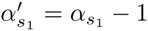; for all the other strategies *s*, 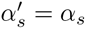.
- For all 1 < s < *n*, 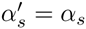.

In the first case, the selected set is broken; one individual playing strategy *s*_1_ is expelled and one individual with strategy *s*_2_ is incorporated to the set. In order to illustrate this case, we take the transition from (*α*_1_*, α*_2_ …, *α_n_)* to (*α*_1_ *+* 1*,α*_2_ … *,α_n_ –*1) as an example. First, a set consisting of *α_s_* strategy *s* individuals is selected, and then breaks with probability *k*(*α*_1_ *+* 1*,α*_2_ … *,α_n_*). Second, a strategy *n* individual is expelled (with probability *α_n_/m*). Finally, a strategy 1 individual is incorporated (with probability *x_A_*, i.e., the fraction of strategy 1 in the population). Thus the transition probability is 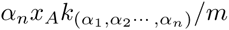. Similarly, the transition probability from state (*α*_1_*, α*_2_ … *,α_n_*) to 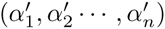, where the two states fulfill the first constraint, is given by

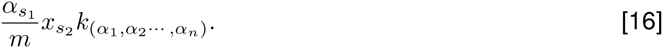

The second case reveals that the two subsequent states are equivalent. Either the selected set is not broken or it is broken but the expelled individual and the new individual are the same in type. In this case, the transition probability can be obtained by the normalisation property of *V* — i.e., one minus the sum of all the other transition probabilities in Eq. (16).

When all the strategies coexist, i.e., 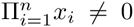, the transition matrix *Q* is aperiodic and irreducible, consequently the Markov chain presents a unique stationary distribution. By Eq. (15), the stationary distribution of *Q* is the same as that of *V*. This holds for any number of total links *l*. However, the size of the state space |*S_n_,_m_*| is 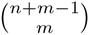 (7). As a consequence, the number of states increases much more rapidly with the set size when there are more than two types of strategies in the population (Fig (2)). Given this, it becomes challenging to calculate the stationary distribution for multiple strategies. Still, as shown in (8), for general *n×n* games and the dynamical network *m* = 2, we have i) that the stationary distribution is a binomial distribution weighted by the duration time, ii) that the conditional transition matrix *V* satisfies the detailed balance condition. This binomial distribution arises from the network structure, which is a special case, *m* = 2, of our set structure. It turns out that these results can be generalised for *m* ≥ 2.

- The stationary distribution of *V, y*, is a multinomial distribution weighted by the duration time, i.e.,

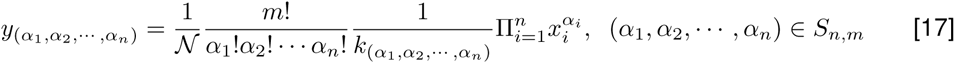

where 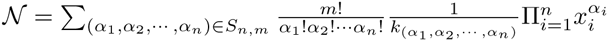 is a normalisation factor.
- The Markov chain *V* fulfills the detailed balance condition, i.e.,

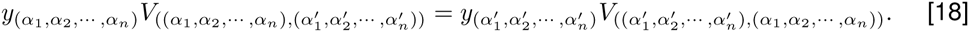 We prove that the distribution Eq. (17) satisfies the detailed balance condition.

If the transition from state (*α*_1;_*α*_2_,…,*α_n_*) to state 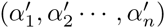 is impossible, then the reverse transition is also impossible. Thus, Eq. (18) holds. In the other cases, the transition is possible. Therefore, the two states (*α*_1;_*α*_2_,…,*α_n_*) and 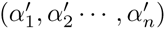 must satisfy one of the two constraints of the transition matrix.

If they fulfill the first constraint, i.e., there exist two different strategies *s*_1_ and *s*_2_ such that 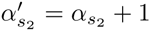 and 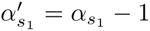; for all the other strategies *s*, 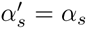. By Eqs. (16) and (17) we have that

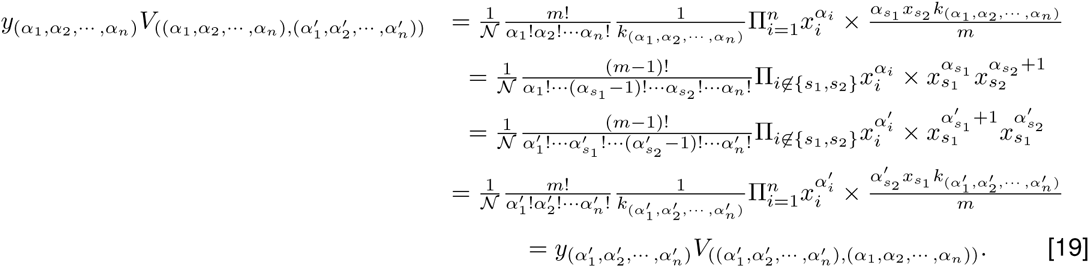

If they fulfill the second constraint, i.e., the two states are the same, then Eq. (18) holds naturally. Therefore the stationary distribution of *Q* is the multinomial distribution weighted by the duration time. Furthermore, *Q* fulfills the detailed balance condition.

When set dynamics are fast, the average payoff is determined by the stationary distribution of each set configuration. For any strategy 1 ≤ *i* ≤ *n*, we have

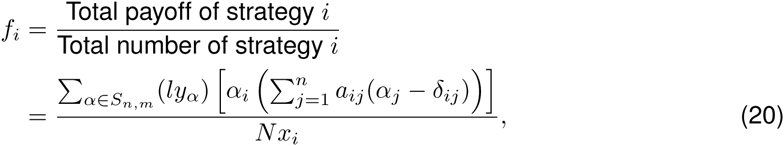

where *δ_ij_* is the Kronecker-delta and *N* is the population size.

Taking Eq. (17) into Eq. (20) leads to

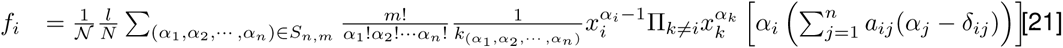

considering that 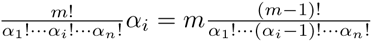 yields that

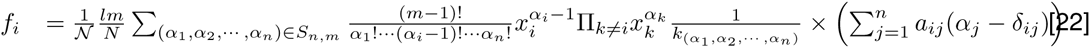

Let 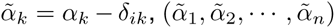 be the co-player configuration of a strategy *i* individual in a set (*α*_1_, *α*_2_,…,*α_n_*). Eq. (22) is given by

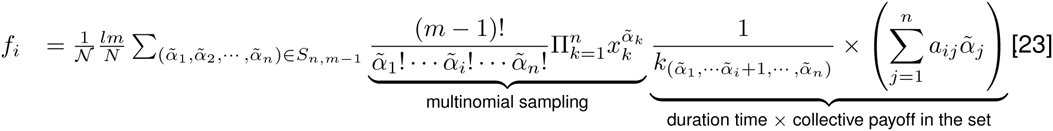

This accumulated payoff is formally equivalent to an *n*-strategy *m*-player game up to a rescaling factor 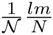. The first term is a multinomial distribution which indicates that *m* – 1 co-players are sampled randomly as if in a well-mixed population. The second term shows that the payoff of strategy *i* of the multi-player game is the collective payoff of strategy *i* in a set times the average duration time of the corresponding set. This term is dependent on (i) the pairwise interaction between strategy *i* and (ii) the set duration time. This explicitly generates an *n*-strategy *m*-player game from a pairwise *n × n* game (*a_ij_*).

#### 3.2 Evolutionary dynamics of strategies

During the imitation process, the role model and the focal individual are both chosen randomly through the entire population. The evolution of strategies can also be approximated by the Langevin Equation. More precisely, in this case when the population is large enough, the demographic noise induced by the finite population size can be neglected (9). This results in the following replicator equation:

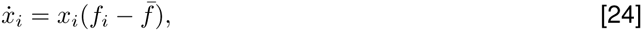

where *f_i_* is given by Eq. (23) and 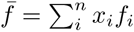 is the average payoff. Consequently, the replicator equation is consistent with an *n*-strategy *m*-player game and can exhibit up to (*n* – 1)*^m^*^−1^ internal isolated equilibria (2).

---

**Figure 1:**
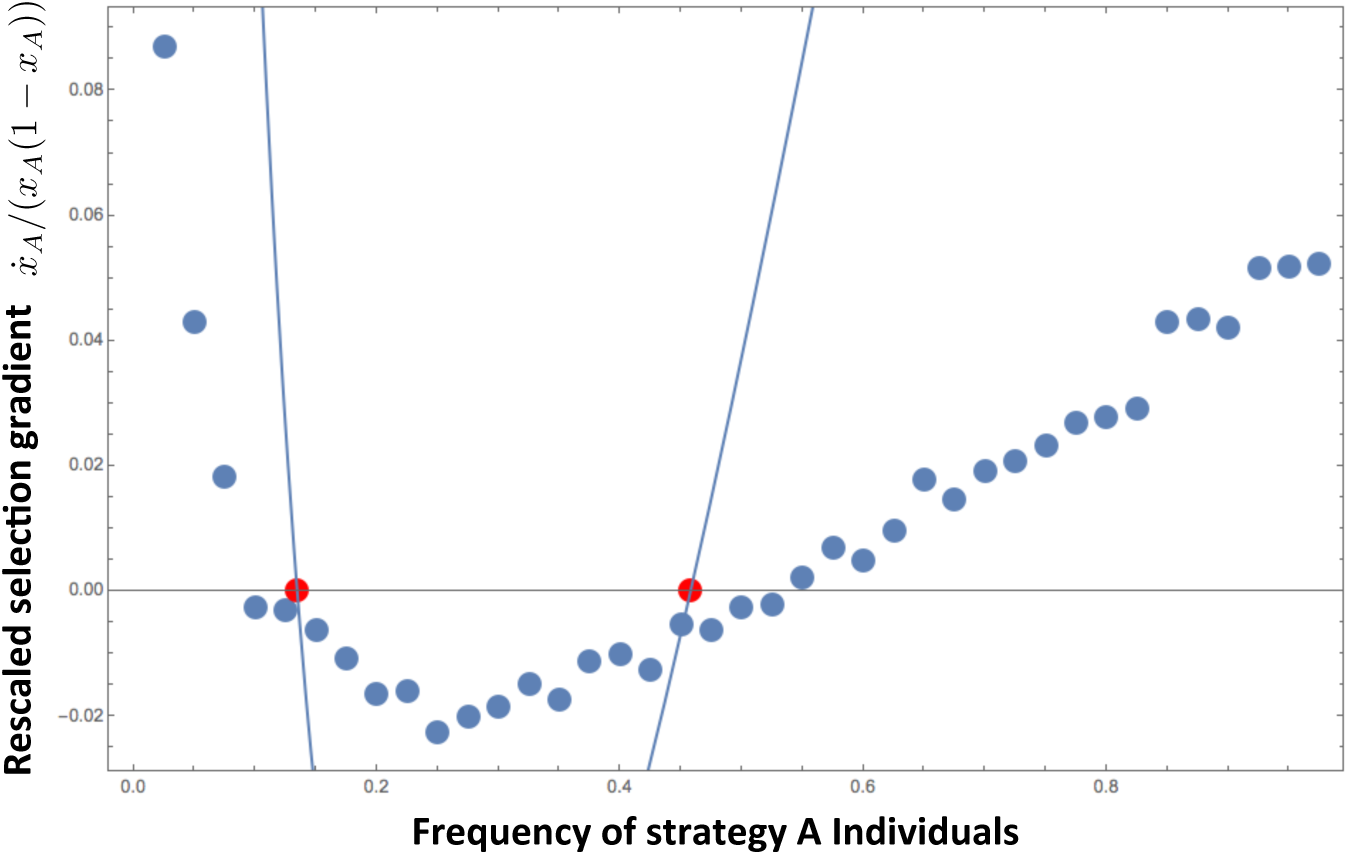
Rescaled selection gradient 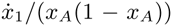. The simulation based on the Fermi update rule (dots) shows that it has two roots. The analytical approximation (Eq.(10)) captures the equilibria of the selection gradient by simulation 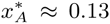 and 0.46 (red dots). The absolute value of the selection gradient, however, is systematically overestimated by the theoretical approximation for the positive selection gradient. This is because there is a heterogeneity in payoffs within the population using the same strategy. Let us assume that 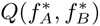 is the probability that a strategy *A* individual is of payoff 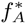 and a strategy *B* individual is of payoff 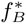. Then the selection gradient based on simulation is an estimator of 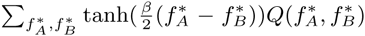. Since tanh(*x*) is convex for *x* > 0, thus the theoretical approximation 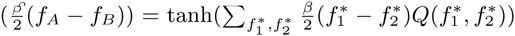 is greater than the estimator of the simulation 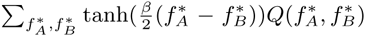. By similar arguments, we obtain that the theoretical approximation underestimates the simulation result for negative selection gradient. Each blue dot in the plot is the average of 100 independent realisations. Every realisation takes 10^7^ generation. For the first 10^4^ generations of each realisation, only set dynamics occur. After that, with a probability of *w* = 10^−3^ two individuals are chosen randomly from the entire population. We keep track of the transition without implementing them. We denote *y* and *z* as the number of times that an individual playing strategy *A* and *B* changes its strategy. 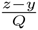 is the estimator of the selection gradient 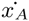, where *Q* is the number of strategy updating events in this realisation. (Parameters: Stag-Hunt game with *a_AA_* = 2, *a_AB_* = 1, *a_BA_* = 1.5 and *a_BB_* = 7. Population size, *N* = 500, number of sets, *l* = 1000, probability of a strategy update, *w* = 10^−3^. Selection intensity, *β* = 0.1. The breaking probabilities are *k_i_* = (1+10*i*)^−1^, where *i* is the number of strategy *A* individuals in the set.)

**Figure 2:**
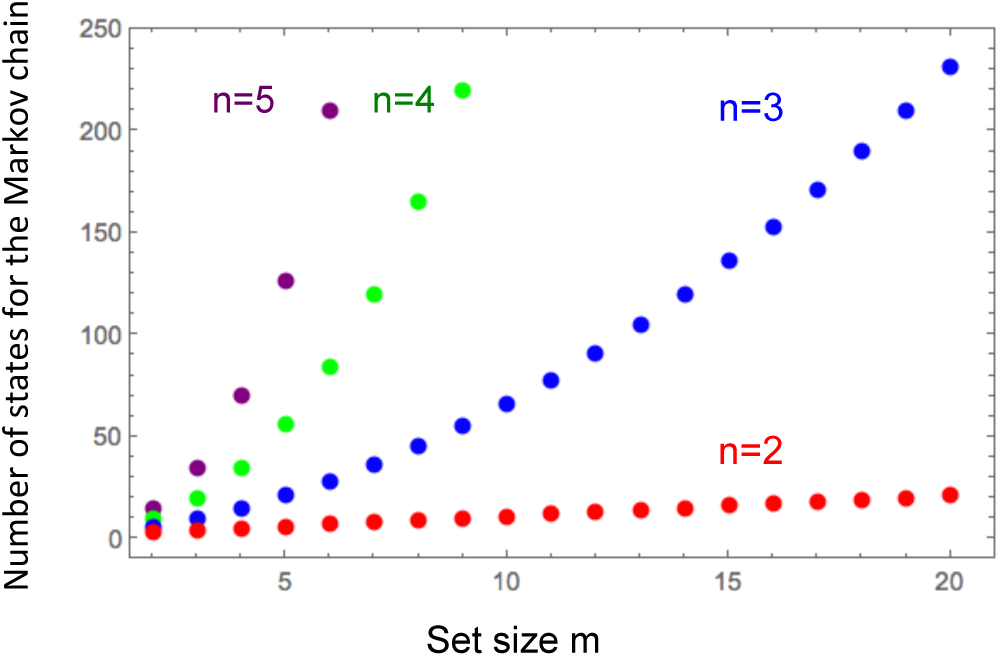
The size of the state space of the Markov chain of the set dynamics as a function of the set size *m*. For a two-strategy game, there are *m* + 1 set configurations. For a three-strategy game, there are 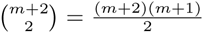 set configurations. In general, the number of the states increases rapidly with the size of the set, if the strategy number increases.

